# Association of five SNPs with human hair colour in the Polish population

**DOI:** 10.1101/087429

**Authors:** A. Siewierska-Górska, A. Sitek, E. Żądzińska, G. Bartosz, D. Strapagiel

## Abstract

Twenty-two variants of the genes involved in hair pigmentation (OCA2, HERC2, MC1R, SLC24A5, SLC45A2, TPCN2, TYR, TYRP1) were genotyped in a group of 186 Polish subjects, representing a range of hair colours (45 red, 64 blond, 77 dark). A genotype-phenotype association analysis was performed.

Using z-statistics and the associated p-value, we identified three variants highly associated with different hair colour categories (rs12913832:A>G in HERC2, rs1805007:T>C and rs1805008:C>T in MC1R). Two variants: rs1800401:C>T in OCA2 and rs16891982:C>G in SLC45A2 showed a high probability of a relation with hair colour, although that probability did not exceed the threshold of statistical significance after applying the Bonferroni correction. We created and validated mathematical logistic regression models in order to test the usefulness of the sets of polymorphisms for hair colour prediction in the Polish population. We subjected four models to stratified cross-validation. The first model consisted of three polymorphisms that proved to be important in the associative analysis. The second model included, apart from the mentioned polymorphisms, additionally rs16891982:C>G in SLC45A. The third model included, apart from the variants relevant in the associative analysis, rs1800401:C>T in OCA. The fourth model consisted of the set of polymorphisms from the first model supplemented with rs16891982:C>G in SLC45A and rs1800401:C>T in OCA. The validation of our models showed that the inclusion of rs16891982:C>G in SLC45A and rs1800401:C>T in OCA increases the prediction of red hair in comparison with the algorithm only including rs12913832:A>G in HERC2, rs1805007:T>C and rs1805008:C>T in MC1R. The model consisting of all the five above-mentioned genetic variants showed good prediction accuracies, expressed by the area under the curve (AUC) of the receiver operating characteristics: 0.84 for the red-haired, 0.82 for the dark-haired and 0.71 for the blond-haired. A genotype-phenotype association analysis brought results similar to those in other studies and confirmed the role of rs16891982:C>G, rs12913832:A>G, rs1805007:T>C and rs1805008:C>T in hair colour determination in the Polish population. Our study demonstrated for the first time the possibility of a share of the rs1800401:C>T SNP in the OCA2 gene in hair colour determination. Including this SNP in the actual hair colour predicting models would improve their predictive accuracy.

## Introduction

Natural pigmentation is a variable human trait that functions mainly as a protection system against the damaging effects of ultraviolet radiation (UVR) (Jablonski and Chaplin, 2000). The pigmentary phenotype is distinctive of geographical ancestry and varies between populations (Shriver et al., 2003). In Europeans, hair colour is highly variable and ranges from light blond to black with an additional variety of red hair shades. Hair colour is a polygenic trait determined by a large subset of genes. These genes encode proteins that either regulate melanin synthesis or act at different stages of it. Melanin is a polymeric pigment produced in the melanosomes, responsible for the colour of the skin, iris and hair (Barsh, 1996; Ortonne and Prota, 1993; Sturm et al., 2003). The presence of different hair colour phenotypes among humans is due to polymorphic substitutions in the pigmentary genes. These single nucleotide polymorphisms (SNPs) lead to subtle changes in the expression or the function of the coded proteins, which in turn affect the production and distribution of melanin subtypes, i.e. eumelanin and pheomelanin. The observed colour is the result of the overall amount of pigment and of the ratio of eumelanin to pheomelanin, which depends on the activity of the rate-limiting enzyme tyrosinase and on the availability of its main substrates: tyrosine and cysteine. The whole pathway is regulated by the melanocortin hormone (αMSH), which acts via its receptor – melanocortin 1 receptor (MC1R) localised in the melanocyte plasma membrane (Ortonne and Prota, 1993; Valverde et al., 1995). It is well known that certain variants of the MC1R gene cause diminished signal transduction, which can result in red hair colour (Beaumont et al., 2007; Schiöth et al., 1999). Although the MC1R and TYR genes seem to play a major role in pigment production, there are other proteins whose function in this process is not fully explained, but their genetic association with phenotype is undeniable (Barsh, 1996). The SLC45A2, TPCN2, SLC24A5 and OCA2 genes encode for the most part transporter proteins that are speculated to supply substrates and cofactors for the melanin synthesis and regulate the pH and the ionic composition of the melanosomal lumen (Brilliant, 2001; Cook et al., 2009; Graf et al., 2005; Lamason et al., 2005; Sulem et al., 2008; Valenzuela et al., 2010). Their allelic variants seem to act as a switch between dark and light hair colour (Cook et al., 2009; Mengel-From et al., 2009). Although huge progress has been made in the field of genetics of human pigmentation and a few genotype-based hair colour determination models for Europeans have been published (Branicki et al., 2011; Kastelic and Drobnič, 2012; Sulem et al., 2007; Valenzuela et al., 2010; Walsh et al., 2013), this topic is still being studied extensively. Many questions regarding the interactions of the genes involved in the process of melanogenesis and the patterns of phenotype inheritance remain to be answered. One must also remember that hair pigmentation is additionally influenced by environmental factors such as sun exposure (Hessefort et al., 2008), diet (Finner, 2013), hormonal metabolism (Slominski et al., 2004; Tobin, 2008; van Beek et al., 2008), and also by an individual’s age (Sitek et al., 2012; Sitek et al., 2013). It is still necessary to continue research in this field, since populations from different geographical regions and of mixed ancestry might differ in the SNP pool that correlates with a given hair colour.

The main goal of this study was to investigate the relations of SNPs within the selected candidate pigmentary genes with hair colour in the Polish population and to determine their effect on hair colour prediction.

## Materials and Methods

### Subjects

All subjects were adult volunteers at the age of over 18 years living in Poland, who donated saliva samples to the Biobank Lab at the Department of Molecular Biophysics of the University of Lodz. Sample collection and anthropological data acquisition through a questionnaire were performed as previously described (Koszarska et al., 2014; Strapagiel et al., 2016). For this study, 186 subjects were randomly chosen and categorised into three distinct groups according to their declared, self-assessed natural hair colour. The mean age of the volunteers at the time of taking samples of biological material and of the examination was 39.97 +/– 14.24 years. Their age ranged from 18 to 74 years. We established groups representing three major hair colour categories according to the Fischer-Saller scale, i.e. red, blond and dark (Table 1). We evened out the proportions of the individuals in each hair colour category by enlarging the red-haired group in order to increase the ability to reveal associations between the categories and genetic variants. Thus, it should be noted that the frequencies of the studied hair colours do not reflect their real proportions in the Polish population. The frequency of red-haired individuals in the Polish population is about 0.7% (0.9% when including red blond), as calculated on the basis of data gathered by the Biobank Lab at the University of Lodz (Supplementary Table 1). The established red-haired group consisted of 45 individuals representing shades of red from blond-red to copper-red. The dark-haired group included 77 people whose hair ranged from light brown through chestnut to black. 64 subjects represented the blond-haired group with shades ranging from platinum blond through blond to dark/honey blond.

**Table 1.** Hair colour classification into three distinct categories

| *Hair colour category* | *Declared hair colour* | *N (%) in the 186 participants group* |
| --- | --- | --- |
| *dark* | *black, dark brown, brown, chestnut* | *77 (41.4)* |
| *blond* | *dark blond, blond, light blond* | *64 (34.4)* |
| *red* | *red, red-blond* | *45 (24.2)* |

### Genotyping

In order to study the association of genetic markers with hair colour, we chose 22 variants only (Table 2, Table 3) within eight different genes involved in the melanin synthesis pathway based on the results of the GWAS – functional and phenotype association studies published through the years (Beaumont et al., 2007; Box et al., 1997; Cook et al., 2009; Flanagan et al., 2000; Graf et al., 2005; Mengel-From et al., 2009; Rebbeck et al., 2002; Sulem et al., 2008; Valenzuela et al., 2010). We selected 22 polymorphisms that were reported as important in determining human phenotypes in European ancestral populations, including hair, skin and iris pigmentation, or in their potential role in skin cancer development (obviously, the chosen polymorphisms do not exhaust all possible SNPs related to the variability of the human pigmentary phenotype).

**Table 2.** Minor allele frequencies (MAF) of the studied variants within three hair colour categories

| *Gene* | *dbSNP* |  | *Allele* | *MAF*<br>*Whole*<br>*sample*<br>*(N=186)* | *MAF*<br>*Red*<br>*(N=45)* | *MAF*<br>*Blond*<br>*(N=64)* | *MAF*<br>*Dark*<br>*(N=77)* |
| --- | --- | --- | --- | --- | --- | --- | --- |
| *HERC2* | *rs12913832* | *NC_000015.10:g.28120472A>G* | A | 0.207 | 0.067 | 0.102 | 0.377 |
| *MC1R* | *rs312262906* | *NC_000016.10:g.89919342_89919343insA* | +A | 0.016 | 0.022 | 0.031 | 0.000 |
| *MC1R* | *rs1805005* | *NC_000016.10:g.89919436G>T* | T | 0.048 | 0.056 | 0.063 | 0.032 |
| *MC1R* | *rs1805006* | *NC_000016.10:g.89919510C>A* | A | 0.008 | 0.000 | 0.016 | 0.006 |
| *MC1R* | *rs34540312* | *NC_000016.10:g.89919523G>C* | C | 0.008 | 0.022 | 0.008 | 0.000 |
| *MC1R* | *rs2228479* | *NC_000016.10:g.89919532G>A* | A | 0.097 | 0.056 | 0.117 | 0.104 |
| *MC1R* | *rs200616835* | *NC_000016.10:g.89919621C>G* | G | 0.003 | 0.011 | 0.000 | 0.000 |
| *MC1R* | *rs11547464* | *NC_000016.10:g.89919683G>A* | A | 0.005 | 0.022 | 0.000 | 0.000 |
| *MC1R* | *rs1805007* | *NC_000016.10:g.89919709C>T* | T | 0.083 | 0.256 | 0.039 | 0.019 |
| *MC1R* | *rs1110400* | *NC_000016.10:g.89919722T>C* | C | 0.024 | 0.022 | 0.000 | 0.045 |
| *MC1R* | *rs1805008* | *NC_000016.10:g.89919736C>T* | T | 0.116 | 0.233 | 0.070 | 0.084 |
| *MC1R* | *rs885479* | *NC_000016.10:g.89919746G>A* | A | 0.022 | 0.000 | 0.039 | 0.019 |
| *MC1R* | *rs1805009* | *NC_000016.10:g.89920138G>C* | C | 0.013 | 0.033 | 0.000 | 0.013 |
| *MC1R* | *rs2228478* | *NC_000016.10:g.89920200A>G* | G | 0.024 | 0.011 | 0.039 | 0.019 |
| *OCA2* | *rs1800401* | *NC_000015.10:g.28014907G>A* | T | 0.051 | 0.011 | 0.031 | 0.091 |
| *SLC24A5* | *rs1426654* | *NC_000015.10:g.48134287A>G* | G | 0.005 | 0.000 | 0.008 | 0.006 |
| *SLC45A2* | *rs16891982* | *NC_000005.10:g.33951588C>G* | G | 0.038 | 0.000 | 0.023 | 0.071 |
| *SLC45A2* | *rs28777* | *NC_000005.10:g.33958854C>A* | C | 0.035 | 0.000 | 0.031 | 0.058 |
| *TPCN2* | *rs35264875* | *NC_000011.10:g.69078931A>T* | T | 0.250 | 0.222 | 0.313 | 0.214 |
| *TPCN2* | *rs3829241* | *NC_000011.10:g.69087895G>A* | A | 0.368 | 0.389 | 0.391 | 0.338 |
| *TYR* | *rs1042602* | *NC_000011.10:g.89178528C>A* | A | 0.347 | 0.344 | 0.391 | 0.312 |
| *TYRP1* | *rs1408799* | *NC_000009.12:g.12672097T>C* | T | 0.304 | 0.233 | 0.328 | 0.325 |

+A – insertion of adenine allele

+A – insertion of adenine allele

**Table 3.** Allelic odds ratios calculated from single SNP hair colour association analysis in the studied sample (N=186)

| Gene | SNP | Allele | Red vs. non-red |  |  |  | Blond vs. non-blond |  |  |  | Dark vs. non-dark |  |  |  |
| --- | --- | --- | --- | --- | --- | --- | --- | --- | --- | --- | --- | --- | --- | --- |
|  |  |  | OR | 95%CI | p | PA | OR | 95%CI | p | PA | OR | 95%CI | p | PA |
| HERC2 | rs12913832:A>G | A | 0.21 | 0.09-0.51 | **0.0005** | 0.97 | 0.32 | 0.17-0.60 | **0.0005** | 0.97 | 6.33 | 3.57-11.22 | **<0.0001** | 1.00 |
| MC1R | rs312262906:insA | +A | 1.58 | 0.28-8.77 | 0.6012 | 0.13 | 3.90 | 0.71-21.61 | 0.1188 | 0.46 | 0.11 | 0.01-1.89 | 0.1269 | 0.45 |
| MC1R | rs1805005:G>T | T | 1.22 | 0.42-3.51 | 0.7162 | 0.10 | 1.56 | 0.60-4.06 | 0.3616 | 0.23 | 0.53 | 0.18-1.52 | 0.2361 | 0.32 |
| MC1R | rs1805006:C>A | A | 0.44 | 0.02-8.62 | 0.5895 | 0.13 | 3.86 | 0.35-42.95 | 0.2723 | 0.29 | 0.71 | 0.06-7.85 | 0.7769 | 0.09 |
| MC1R | rs34540312:G>C | C | 6.39 | 0.57-71.28 | 0.1320 | 0.44 | 0.95 | 0.09-10.61 | 0.9686 | 0.25 | 0.20 | 0.01-3.89 | 0.2871 | 0.28 |
| MC1R | rs2228479:G>A | A | 0.48 | 0.18-1.26 | 0.1364 | 0.44 | 1.41 | 0.70-2.84 | 0.3365 | 0.25 | 1.15 | 0.57-2.29 | 0.6963 | 0.10 |
| MC1R | rs200616835:C>G | G | 9.47 | 0.38-234.52 | 0.1698 | 0.39 | 0.63 | 0.03-15.62 | 0.7789 | 0.09 | 0.47 | 0.02-11.60 | 0.6438 | 0.12 |
| MC1R | rs11547464:G>A | A | 15.96 | 0.76-335.60 | 0.0747 | 0.55 | 0.38 | 0.02-7.92 | 0.5304 | 0.20 | 0.28 | 0.01-5.88 | 0.4127 | 0.15 |
| MC1R | rs1805007:C>T | T | 11.76 | 5.04-27.44 | **<0.0001** | 1.00 | 0.34 | 0.13-0.91 | 0.0317 | 0.69 | 0.13 | 0.04-0.45 | 0.0012 | 0.94 |
| MC1R | rs1110400:T>C | C | 0.89 | 0.18-4.38 | 0.8889 | 0.07 | 0.10 | 0.01-1.67 | 0.1080 | 0.48 | 5.14 | 1.05-25.10 | 0.0429 | 0.65 |
| MC1R | rs1805008:C>T | T | 3.60 | 1.90-6.92 | **0.0001** | 0.99 | 0.47 | 0.22-1.01 | 0.0522 | 0.61 | 0.58 | 0.29-1.15 | 0.1173 | 0.47 |
| MC1R | rs885479:G>A | A | 0.18 | 0.01-3.12 | 0.2379 | 0.32 | 3.27 | 0.77-13.89 | 0.1091 | 0.48 | 0.85 | 0.20-3.60 | 0.8212 | 0.08 |
| MC1R | rs1805009:G>C | C | 4.83 | 0.79-29.36 | 0.0874 | 0.52 | 0.17 | 0.01-3.09 | 0.2307 | 0.33 | 0.94 | 0.16-5.71 | 0.9491 | 0.06 |
| MC1R | rs2228478:A>G | G | 0.38 | 0.05-3.12 | 0.3711 | 0.22 | 2.44 | 0.64-9.25 | 0.1897 | 0.34 | 0.70 | 0.17-2.85 | 0.6207 | 0.12 |
| OCA2 | rs1800401:C>T | T | 0.16 | 0.02-1.25 | 0.0814 | 0.54 | 0.49 | 0.16-1.52 | 0.2169 | 0.34 | 4.26 | 1.50-12.09 | 0.0065 | 0.86 |
| SLC24A5 | rs1426654:A>G | G | 0.62 | 0.03-13.03 | 0.7583 | 0.09 | 1.91 | 0.12-30.85 | 0.6473 | 0.12 | 1.42 | 0.09-22.85 | 0.8054 | 0.08 |
| SLC45A2 | rs16891982:C>G | G | 0.10 | 0.01-1.73 | 0.1143 | 0.47 | 0.51 | 0.14-1.86 | 0.3058 | 0.37 | 5.51 | 1.51-20.11 | 0.0097 | **0.83** |
| SLC45A2 | rs28777:C>A | C | 0.11 | 0.01-1.87 | 0.1272 | 0.45 | 0.84 | 0.25-2.79 | 0.7788 | 0.09 | 3.32 | 1.00-10.99 | 0.0493 | 0.62 |
| TPCN2 | rs35264875:A>T | T | 0.82 | 0.47-1.44 | 0.4850 | 0.17 | 1.64 | 1.01-2.65 | 0.0447 | 0.64 | 0.72 | 0.44-1.17 | 0.1821 | 0.38 |
| TPCN2 | rs3829241:G>A | A | 1.12 | 0.69-1.83 | 0.6416 | 0.12 | 1.16 | 0.74-1.80 | 0.5177 | 0.16 | 0.80 | 0.52-1.23 | 0.3038 | 0.27 |
| TYR | rs1042602:C>A | A | 0.99 | 0.60-1.63 | 0.9575 | 0.06 | 1.34 | 0.86-2.09 | 0.1987 | 0.36 | 0.77 | 0.49-1.19 | 0.2326 | 0.32 |
| TYRP1 | rs1408799:T>C | T | 0.63 | 0.36-1.09 | 0.0969 | 0.50 | 1.19 | 0.75-1.89 | 0.4595 | 0.18 | 1.18 | 0.76-1.85 | 0.4612 | 0.18 |

+A – insertion of adenine; OR – odds ratio; 95%CI – 95% confidence interval for OR;

+A – insertion of adenine; OR – odds ratio; 95%Cl – 95% confidence interval for OR; p – value associated with z-statistics (https://www.medcalc.net/tests/odds_ratio.php); statistically significant results (after applying the Benferroni correction p<0,0008) are in bold; PA-power analysis (1-β),

The isolation of the genomic DNA from saliva was performed as previously reported (Koszarska et al., 2014). Genotyping of the variants within the studied genes (Table 2) was performed with the HRM (High Resolution Melt) method accompanied by direct sequencing. Genomic DNA of the subjects was used as a template for PCR amplification of the target region containing the SNP of interest with a set of primers designed by the authors (Supplementary Table 2) using Primer3 software (Koressaar and Remm, 2007; Untergasser et al., 2012). The details of the HRM method employed were described previously (Koszarska et al., 2014). We used direct sequencing as an HRM genotyping reference method in order to confirm the sample cluster allele genotype or as an alternative method in the case of the MC1R gene, for which the whole exon was sequenced using two overlapping primer pairs. Supplementary Table 3 presents the set of primers used for the amplification, all designed and verified by the authors of this article using Primer3 software. For the purpose of sequencing, we amplified DNA fragments with PCR using 1 ng of genomic DNA as a template, 2xGoTaq Green polymerase master mix (Promega) and the above-mentioned primers (Supplementary Table 3). The samples were thermocycled in the Bio-Rad Thermal Cycler-1000 (Bio-Rad Laboratories Inc., Hercules, CA, USA). Prior to sequencing, all fragments obtained by PCR amplification were visualised using 1% agarose gels and purified by Wizard^®^ SV Gel and PCR Clean-Up System (Promega Corporation, Madison, WI, USA) according to the manufacturer’s protocol. Direct sequencing was performed by outside companies: Eurofins MWG Operon (Ebersberg, Germany) or Genomed S.A. (Warsaw, Poland). The results were analysed using the Codon Code Aligner program (CodonCode™ Aligner-Software (CodonCode Corporation, Centerville, MA, USA)).

### Statistical Analysis

For each of the studied variants, minor allele frequencies (MAF) were calculated within each hair colour category, and the Hardy-Weinberg equilibrium was estimated. We evaluated the association between each allele of the verified genotype and of the hair colour phenotype using z-statistics and the associated p-value (https://www.medcalc.net/tests/odds_ratio.php). When analysing the statistical significance of the differences, the Bonferroni correction for multiple testing was applied, and differences with p<0.0008 were considered statistically significant (Table 3). Having identified the polymorphisms, which had statistically significantly differentiated the isolated groups by means of multivariable logistic regression (STATISTICA software version 10.0, StatSoft Inc. Poland), we checked to what extent those SNPs enabled hair colour prediction in the studied group. Additionally, we assessed the share of two polymorphisms in hair colour prediction, whose probability of a relation with this trait was the highest, but did not exceed the threshold of statistical significance after applying the Bonferroni correction.

## Results

### Genotype-phenotype association

All the genetic variants investigated in this study together with their descriptions and genomic reference sequence accession numbers (RefSeq) are presented in Table 2. Further in the text, while describing the changes in mRNA and their predicted protein variants, the following reference sequences were used: NM_002386.3 (MC1R), NM_000275.2 (OCA2), NM_205850.2 (SLC24A5), NM_001012509.2 (SLC45A2), NM_139075.3 (TPCN2), and NM_000372.4 (TYR). No statistically significant departures from the Hardy-Weinberg equilibrium were detected for the studied SNPs. Due to the moderate sample size, we decided to divide the volunteers into only three hair colour categories in order to increase statistical reliability. Genome data were deposited at the European Genome-phenome Archive (EGA), which is hosted at the EBI and the CRG, under accession number EGAS00001000997. Minor allele frequencies were calculated for each variant both in the whole studied sample and in each hair colour category (Table 2). After the genotype-phenotype association analysis, we identified three variants highly associated with different hair colour categories (rs12913832:A>G in HERC2, rs1805007:T>C and rs1805008:C>T in MC1R) (Table 3). MC1R proved to be the most significant gene in red hair colour determination. c.451C>T (p.(Arg151Cys)) and c.478C>T (p.(Arg160Trp)) substitutions acted as the strongest predictors for this hair colour category with OR=11.76 and OR=3.60, respectively. Not only did the rs1805007:T>C variants account for red hair, but they also showed a substantial relation with non-red hair colour, scoring an OR of 0.13 for dark hair (95% CI: [0.04-0.45]; p=0.0012) (Table 3). Our results showed rs12913832:A>G in HERC2 to be the strongest indicator of dark hair among all the studied variants (OR=6.33; 95% CI: [3.57-11.22]; p<0.0001). This SNP also showed a negative correlation with non-dark hair colour categories manifesting an OR of 0.21 for the red-haired group (CI 95% [0.09-0.51]; p=0.0005) and an OR of 0.32 for the blond-haired group (CI 95% [0.17-0.60]; p=0.0005) (Table 3). We also found that two polymorphisms showed a high probability of a relation with dark hair: rs1800401:C>T (OR=4.26; CI 95% [1.50 – 12.09]; p=0.0065) and rs16891982:C>G (OR=5.51; CI 95% [1.51 – 20.11]; p=0.0097], although that probability did not exceed the threshold of statistical significance after applying the Bonferroni correction (Table 3).

### Modelling of the prediction tool for human hair colour pigmentation – verification of the observed associations

Based on the results of the associative analysis, we constructed hair colour prediction models, using logistic regression. Four models were tested. The first model consisted of three polymorphisms that proved to be important in the associative analysis. The second model included, apart from the mentioned polymorphisms, additionally rs16891982:C>G in SLC45A. The third model included, apart from the variants relevant in the associative analysis, rs1800401:C>T in OCA. The fourth model consisted of the set of polymorphisms from the first model supplemented with rs16891982:C>G in SLC45A and rs1800401:C>T in OCA.

An individual’s hair colour (y) was predicted by the number of the minor alleles in the genotype of each SNP (0, 1, and 2). In order to assess the predictive usefulness of the individual sets of SNPs, stratified cross-validation was performed, i.e. such a division of the objects between the training set and the test set that the original proportions between the decision classes were preserved. For this purpose, the entire studied population was divided into 10 separable subpopulations: 9 with 18 subjects in each (8 dark, 4 red and 6 blond) and 1 with 24 subjects (5 dark, 9 red and 10 blond). Each of the 10 subpopulations was regarded to be a test subset, while the other 9 subpopulations made a training set.

We used the AUC of the ROC curve as a universal measure of the prediction accuracy of our model along with positive predictive value (PPV), negative predictive value (NPV), accuracy (ACC), and sensitivity and specificity parameters at a fixed threshold of 0.5.

The mean results of the tenfold cross-validation showed that separate inclusion of each of the tested variants of the SLC45A2 and OCA2 genes in the initial model (comprising polymorphisms statistically significantly related to hair colour), slightly improved the prediction of red hair (an increase of the AUC by 0.017 for rs16891982:C>G in SLC45A2 and by 0.011 for rs1800401:C>T in OCA2). The inclusion of each of these genetic variants also reduced the standard deviation of the field under the ROC curve in comparison with the initial model. However, the greatest change was observed in the situation when both variants of the SLC45A2 and OCA2 genes were included in the first model (an increase of the AUC by 0.028 for the red-haired and a reduction of the SD AUC by 0.018) (Table 4).

**Table 4.** Hair colour prediction accuracy of the logistic regression model

|  | *Model 1* |  |  | *Model 2* |  |  | *Model 3* |  |  | *Model 4* |  |  |
| --- | --- | --- | --- | --- | --- | --- | --- | --- | --- | --- | --- | --- |
|  | *model built with 3 SNPs*<br>*(rs1805007:C>T,*<br>*rs1805008:C>T,*<br>*rs12913832:A>G)* |  |  | *model built with 4 SNPs*<br>*(rs1805007:C>T,*<br>*rs1805008:C>T,*<br>*rs12913832:A>G*<br>***rs16891982:C>G)*** |  |  | *model built with 4 SNPs*<br>*(rs1805007:C>T,*<br>*rs1805008:C>T,*<br>*rs12913832:A>G,*<br>***rs1800401:C>T)*** |  |  | *model built with 5 SNPs*<br>*(rs1805007:C>T,*<br>*rs1805008:C>T,*<br>*rs12913832:A>G,*<br>***rs16891982:C>G,***<br>***rs1800401:C>T)*** |  |  |
| *averaged*<br>*results for*<br>*the 10*<br>*testing*<br>*sets* | *Red* | *Blond* | *Dark* | *Red* | *Blond* | *Dark* | *Red* | *Blond* | *Dark* | *Red* | *Blond* | *Dark* |
| *AUC* | 0.809 | 0.708 | 0.821 | 0.826 | 0.703 | 0.822 | 0.820 | 0.707 | 0.817 | 0.837 | 0.706 | 0.824 |
| *SD AUC* | 0.110 | 0.097 | 0.078 | 0.098 | 0.114 | 0.093 | 0.096 | 0.106 | 0.076 | 0.092 | 0.125 | 0.088 |
| *Sensitivity*<br>*[%]* | 51.67 | 66.00 | 61.75 | 51.67 | 62.67 | 68.00 | 55.56 | 64.33 | 65.50 | 66.67 | 61.00 | 68.00 |
| *Specificity*<br>*[%]* | 95.71 | 70.12 | 88.37 | 95.71 | 73.45 | 85.37 | 95.00 | 72.62 | 87.37 | 92.86 | 75.95 | 84.37 |
| *PPV [%]* | 73.33 | 56.58 | 78.52 | 73.33 | 57.59 | 76.58 | 74.05 | 58.13 | 78.16 | 74.17 | 59.64 | 75.19 |
| *NPV [%]* | 86.91 | 79.30 | 75.98 | 86.91 | 78.51 | 78.48 | 87.90 | 79.21 | 77.73 | 90.40 | 78.36 | 78.42 |
| *ACC [%]* | 85.42 | 68.47 | 76.39 | 85.42 | 69.58 | 77.50 | 85.56 | 69.58 | 77.50 | 86.53 | 70.69 | 76.94 |

AUC – area under ROC curve; SD AUC – standard deviation for AUC; PPV – positive predictive value; NPV – negative predictive value; ACC – prediction accuracy

The sensitivity, the specificity, the PPV, the NPV and the ACC did not change after the inclusion of rs16891982:C>G in SLC45A2 in comparison with the initial model, but their values were increased by the inclusion of rs1800401:C>T in the first model (model 4). All these characteristics were best improved by the inclusion of both the above-mentioned SNPs in the model (model 5): the sensitivity of prediction of red hair increased by 15 percentage points with a relatively small decrease of specificity (by 2.85 percentage points), the PPV increased by 0.84 percentage points, the NPV increased by 3.49 percentage points, and the ACC increased by 1.11 percentage points (Table 4).

Rs16891982:C>G in SLC45A2 and rs1800401:C>T in OCA2 did not change the predictive quality of the initial model for blond and dark hair (Table 4).

## Discussion

### Polymorphisms significant in the prediction of the hair colour phenotype

The hair colour association analysis of the studied SNPs revealed that three variants significantly contributed to hair colour determination in Polish population: rs12913832:A>G in HERC2, rs1805007:T>C and rs1805008:C>T in MC1R. MC1R proved to be the most significant gene in red hair colour determination. These observations are in accordance with previous studies of the MC1R gene’s involvement in pigmentary phenotype variation where the two variants also greatly contributed to red hair colour determination regardless of the geographical ancestry of the studied population (Beaumont et al., 2007; Box et al., 1997; Branicki et al., 2007; Flanagan et al., 2000; Sulem et al., 2007; Valenzuela et al., 2010). There was a number of other SNPs within MC1R that seemed distinctive of the red-haired group, as their penetrance for this colour category reached 100% in our study (rs200616835:C>G, rs11547464:G>A). However, these variants did not prove to be statistically relevant in the genotype-phenotype association, probably due to the very low frequency of their minor alleles combined with the relatively small size of the red-haired sample. Interestingly, there were also MC1R alleles with 0% presence in the red-haired group, since they were identified only in the non-red individuals: c.252C>A (p.(Asp84Glu)) (rs1805006) and c.488G>A (p.(Arg163Gln)) (rs885479) (Table 2). As regards c.488G>A (p.(Arg163Gln)), other researchers (Box et al., 1997; Flanagan et al., 2000) had similar observations, considered it to be correlated rather with geographical and ethnic descent than with visible pigmentary traits and generally designated it as an *r* allele of low (Branicki et al., 2007; Branicki et al., 2011) or of no significance (Mengel-From et al., 2009). The c.252C>A (p.(Asp84Glu)) substitution is somewhat controversial, since it was previously reported to diminish the receptor’s functional ability (Beaumont et al., 2007) and to be characteristic of red-haired people (Branicki et al., 2007; Valverde et al., 1995). What supports our lack of association of this allele is the fact that we only found a heterozygous state in the checked subjects, so we were not able to predict what the hair phenotype could look like in variant homozygous people. However, Flanagan et al. (Flanagan et al., 2000) reported this SNP to be one present both in red and in non-red-haired individuals. Box et al. (Box et al., 1997) did not find any association with hair colour for this variant, either; however, they observed its incidence in the red-haired group. A rather substantial discrepancy with previous studies arose for the rs18005009, c.800G>C (p.(Asp294His)) variant, which we did not find significantly associated (OR=4.83, but p=0.0874; Table 3) with red hair colour, although it has been indicated by other authors to be a strong marker of red hair (Beaumont et al., 2007; Box et al., 1997; Branicki et al., 2007; Branicki et al., 2011; Flanagan et al., 2000; Valverde et al., 1995).

Our results show rs12913832:A>G in HERC2 to be the strongest indicator of dark hair among all the studied variants. Similar observations had been previously made by other researchers (Branicki et al., 2011; Cook et al., 2009; Valenzuela et al., 2010). Lately, we demonstrated a relation of this SNP with dark hair colour in Polish children at the age of 7-10 years (Sitek et al., 2016). Some authors (Mengel-From et al., 2009) also observed that this SNP contributed to dark pigmentation of the iris. A direct association of just this single variant with blond hair might be caused by the fact that this is an intermediate hair colour category that has a tendency to change in an individual during his/her ontogeny. Many people who are blond during childhood or early adulthood switch to darker hair colour later in life (Rees, 2003; Sitek et al., 2013). It would be a good idea to compare the genetic makeup of people who had blond hair consistently throughout their entire life with those who became darker-haired after adolescence. This kind of study would allow us to find out which variants truly contribute to blond hair colour.

Previously, the OCA2 gene was predominantly referred to in various studies of the genetic determinants of iris colour (Duffy et al., 2007; Rebbeck et al., 2002). In the genotype-phenotype association study, we found that rs1800401:C>T was characterised by a high probability of a relation with dark hair, although that probability did not exceed the threshold of statistical significance after applying the Bonferroni correction. We attempted to prove the earlier hypothesis by Rebbeck et al. (Rebbeck et al., 2002), who were the first to indicate a link between this SNP and hair coloration, but fell short of statistical proof. Our latest research points to a share of this SNP in the determination of dark hair in Polish children at the prepubertal age (Sitek et al., 2016). Interestingly, this variant has been previously tested for hair colour association in a Queensland population (Duffy et al., 2007), but their haplotype association study did not provide a clear answer to the question whether this SNP contributed to hair colour. It was also tested in a Polish population before, but no association with hair colour was found (Branicki et al., 2011). Also in a Danish-Scottish population, it did not show any statistical significance, although the MAF value and the sample size were similar to our sample group (Mengel-From et al., 2009). Our finding is also in disagreement with the results of Valenzuela et al. (Valenzuela et al., 2010), who tested the correlation of several SNPs with total melanin production and distribution in the hair, iris and skin in a population from Arizona (USA). These authors did not find any association of this variant with melanin content and melanin ratio, but reported that several other SNPs within OCA2 were involved in melanin content regulation. Our data do not allow us to put forward any simple hypothesis about where these discrepancies come from.

Based on our association results, we can conclude that one variant in the SLC45A2 gene (rs16891982:C>G) is characterised by a relatively high probability of a relation with dark hair colour, although the degree of this probability did not exceed the threshold of statistical significance after applying the Bonferroni correction. Probably, also zero penetrance of its minor alleles in the red-haired group contributes to our hair colour prediction accuracy. Mengel-From et al. (Mengel-From et al., 2009) calculated an OR of 7.0 for the association of rs16891982:C>G with dark hair in the Danish population, which is very close to our results. On the other hand, in a study of Southern European population by Cook et al. (Cook et al., 2009), no association between this SNP and hair or iris colour was found, although a relation with darker skin tone was indicated. This functional study reported higher melanin levels and elevated tyrosinase activity in melanocyte strains carrying this particular variant, which may underlie a relation of this SNP with pigmentary traits. Our earlier research conducted on 7-10-year-old Polish children confirmed a relation of rs16891982:C>G with high values of the spectrophotometrically assessed skin melanin index, but it did not demonstrate its association with hair colour determined by means of the Fischer-Saller scale (Sitek et al., 2016).

Consistently with the data published by Branicki et al. (Branicki et al., 2011), rs1426654:A>G in SLC24A5 (MAF=0.005; Table 2) had no significant association with hair pigmentation in the Polish population. This SNP had a very low frequency of the minor alleles in our studied group, which was also noted by other authors (Branicki et al., 2011; Lamason et al., 2005) in populations of European descent. This locus has been evaluated as Ancestry Informative Region (Draus-Barini et al., 2013). On the contrary, Valenzuela et al. (Valenzuela et al., 2010) reported this variant to be an informative one, taking into account hair melanin variance in individuals of various ethnic backgrounds, and used it as one of three markers in a total hair melanin multiple linear regression model. Similarly, the functional study by Cook et al. (Cook et al., 2009), reported that this variant was responsible for elevated levels of melanin content and tyrosinase activity in the cultured melanocytes, but the complementary genotype-phenotype study did not reveal its relevant association with the pigmentary phenotype.

### Modelling of the prediction tool for human hair colour pigmentation – verification of the observed associations

In comparison with the prediction model for the Polish population published by Branicki et al. (Branicki et al., 2011), our algorithms had slightly lower prediction accuracies for red and blond hair and slightly higher prediction accuracies for dark hair. The model of Branicki et al. (Branicki et al., 2011) scored AUC values of 0.75 for blond, 0.72 for brown, 0.90 for red and 0.78 for black hair in the validation dataset. It should be noted that the model of Branicki et al. (Branicki et al., 2011) was built on the basis of 385 volunteers, while the model in our study is only based on 186 individuals, which could affect the results of prediction. At the same time, the predictive values are calculated for separate hair colour categories, which are only three in our study, whereas Branicki et al. (Branicki et al., 2011) divided hair colour into either four categories (blond, red, brown and black) or into seven categories (blond, dark blond, brown, black, auburn, blond-red and red). It is also worth noting that Branicki et al. (Branicki et al., 2011) had a different approach towards the variants of the MC1R gene, which were pooled into two groups, as established previously in the literature, designated as *R* and *r*. We decided to keep the variants as separate markers, since in our view they provided independent information about an individual’s hair colour. Currently, the most widely used system capable of simultaneous prediction of hair and eye colour is HIrisPlex. The HIrisPlex model average prediction accuracies for four hair colour categories were as follows: 69.5% for blond, 78.5% for brown, 80% for red and 87.5% for black hair (Walsh et al., 2013).

In conclusion, after the genotype-phenotype association analysis, we identified three variants highly associated with different hair colour categories in the Polish population: rs12913832:A>G in HERC2, rs1805007:T>C and rs1805008:C>T in MC1R. Rs1800401:C>T in OCA2 and rs16891982:C>G in SLC45A2 were characterised by a high probability of a relation with dark hair colour in adult Poles, although that probability did not exceed the threshold of statistical significance after applying the Bonferroni correction. Rs16891982:C>G in SLC45A2 is a variant with an already recognised share in hair colour prediction (it is part of the HIrisPlex model). Based on the results of cross-validation, we suggest for the first time that incorporating rs1800401:C>T in the existing models for predicting hair colour could increase their accuracy.

### Competing interests

The authors declare that they have no competing interests.

Ethical approval was obtained from the Research Bioethics Commission of the University of Lodz.

## Acknowledgments

We would like to thank sincerely Prof. Wojciech Branicki from the Institute of Forensic Research, Cracow, Poland, for critical reviews of the manuscript. This work was supported by the EU through the European Regional Development Fund (Project POIG 01.01.02-10-005/08). Results obtained during that research were included into patent pending technical report – P.403360.

## Authors’ contribution

DS served as the principal investigator for the research and analysed data; AS performed the statistical analysis and analysed data; ASG designed and carried out the genetic laboratory analysis and interpreted the results, performed the research project and analysed data. EŻ and GB designed the concept of the research and analysed data. All the authors were involved in drafting the manuscript and approved the final manuscript.

Supplementary information is available at the HOMO Journal of Comparative Human Biology website.

## References

Barsh, G.S., 1996. The genetics of pigmentation: from fancy genes to complex traits. Trends Genet 12, 299–305.

Beaumont, K.A., Shekar, S.N., Newton, R.A., James, M.R., Stow, J.L., Duffy, D.L., Sturm, R.A., 2007. Receptor function, dominant negative activity and phenotype correlations for MC1R variant alleles. Hum Mol Genet 16, 2249–2260.

Box, N.F., Wyeth, J.R., O'Gorman, L.E., Martin, N.G., Sturm, R.A., 1997. Characterization of melanocyte stimulating hormone receptor variant alleles in twins with red hair. Hum Mol Genet 6, 1891–1897.

Branicki, W., Brudnik, U., Kupiec, T., Wolańska-Nowak, P., Wojas-Pelc, A., 2007. Determination of phenotype associated SNPs in the MC1R gene. J Forensic Sci 52, 349–354.

Branicki, W., Liu, F., van Duijn, K., Draus-Barini, J., Pośpiech, E., Walsh, S., Kupiec, T., Wojas-Pelc, A., Kayser, M., 2011. Model-based prediction of human hair color using DNA variants. Human Genetics 129, 443–454.

Brilliant, M.H., 2001. The mouse p (pink-eyed dilution) and human P genes, oculocutaneous albinism type 2 (OCA2), and melanosomal pH. Pigment Cell Res 14, 86–93.

Cook, A.L., Chen, W., Thurber, A.E., Smit, D.J., Smith, A.G., Bladen, T.G., Brown, D.L., Duffy, D.L., Pastorino, L., Bianchi-Scarra, G., Leonard, J.H., Stow, J.L., Sturm, R.A., 2009. Analysis of cultured human melanocytes based on polymorphisms within the SLC45A2/MATP., SLC24A5/NCKX5, and OCA2/P loci. J Invest Dermatol 129, 392–405.

Draus-Barini, J., Walsh, S., Pośpiech, E., Kupiec, T., Głąb, H., Branicki, W., Kayser, M., 2013. Bona fide colour: DNA prediction of human eye and hair colour from ancient and contemporary skeletal remains. Investig Genet 4, 3.

Duffy, D.L., Montgomery, G.W., Chen, W., Zhao, Z.Z., Le, L., James, M.R., Hayward, N.K., Martin, N.G., Sturm, R.A., 2007. A three-single-nucleotide polymorphism haplotype in intron 1 of OCA2 explains most human eye-color variation. Am J Hum Genet 80, 241–252.

Finner, A.M., 2013. Nutrition and hair: deficiencies and supplements. Dermatol Clin 31, 167–172.

Flanagan, N., Healy, E., Ray, A., Philips, S., Todd, C., Jackson, I.J., Birch-Machin, M.A., Rees, J.L., 2000. Pleiotropic effects of the melanocortin 1 receptor (MC1R) gene on human pigmentation. Hum Mol Genet 9, 2531–2537.

Graf, J., Hodgson, R., van Daal, A., 2005. Single nucleotide polymorphisms in the MATP gene are associated with normal human pigmentation variation. Hum Mutat 25, 278–284.

Hessefort, Y., Holland, B.T., Cloud, R.W., 2008. True porosity measurement of hair: a new way to study hair damage mechanisms. J Cosmet Sci 59, 303–315.

Jablonski, N.G., Chaplin, G., 2000. The evolution of human skin coloration. J Hum Evol 39, 57–106.

Kastelic, V., Drobnič, K., 2012. A single-nucleotide polymorphism (SNP) multiplex system: the association of five SNPs with human eye and hair color in the Slovenian population and comparison using a Bayesian network and logistic regression model. Croat Med J 53, 401–408.

Koressaar, T., Remm, M., 2007. Enhancements and modifications of primer design program Primer3. Bioinformatics 23, 1289–1291.

Koszarska, M., Kucsma, N., Kiss, K., Varady, G., Gera, M., Antalffy, G., Andrikovics, H., Tordai, A., Studzian, M., Strapagiel, D., Pulaski, L., Tani, Y., Sarkadi, B., Szakacs, G., 2014. Screening the expression of ABCB6 in erythrocytes reveals an unexpectedly high frequency of Lan mutations in healthy individuals. PloS one 9, e111590.

Lamason, R.L., Mohideen, M.A., Mest, J.R., Wong, A.C., Norton, H.L., Aros, M.C., Jurynec, M.J., Mao, X., Humphreville, V.R., Humbert, J.E., Sinha, S., Moore, J.L., Jagadeeswaran, P., Zhao, W., Ning, G., Makalowska, I., McKeigue, P.M., O'Donnell, D., Kittles, R., Parra, E.J., Mangini, N.J., Grunwald, D.J., Shriver, M.D., Canfield, V.A., Cheng, K.C., 2005. SLC24A5, a putative cation exchanger, affects pigmentation in zebrafish and humans. Science 310, 1782–1786.

Mengel-From, J., Wong, T.H., Morling, N., Rees, J.L., Jackson, I.J., 2009. Genetic determinants of hair and eye colours in the Scottish and Danish populations. BMC Genet 10, 88.

Ortonne, J.P., Prota, G., 1993. Hair melanins and hair color: ultrastructural and biochemical aspects. J Invest Dermatol 101, 82S–89S.

Rebbeck, T.R., Kanetsky, P.A., Walker, A.H., Holmes, R., Halpern, A.C., Schuchter, L.M., Elder, D.E., Guerry, D., 2002. P gene as an inherited biomarker of human eye color. Cancer Epidemiol Biomarkers Prev 11, 782–784.

Rees, J.L., 2003. Genetics of hair and skin color. Annu Rev Genet 37, 67–90.

Schiöth, H.B., Phillips, S.R., Rudzish, R., Birch-Machin, M.A., Wikberg, J.E., Rees, J.L., 1999. Loss of function mutations of the human melanocortin 1 receptor are common and are associated with red hair. Biochem Biophys Res Commun 260, 488–491.

Shriver, M.D., Parra, E.J., Dios, S., Bonilla, C., Norton, H., Jovel, C., Pfaff, C., Jones, C., Massac, A., Cameron, N., Baron, A., Jackson, T., Argyropoulos, G., Jin, L., Hoggart, C.J., McKeigue, P.M., Kittles, R.A., 2003. Skin pigmentation, biogeographical ancestry and admixture mapping. Hum Genet 112, 387–399.

Sitek, A., Rosset, I., Żądzińska, E., Siewierska-Górska, A., Pierowska, A., Strapagiel, D., 2016. Selected gene polymorphisms effect on skin and hair pigmentation in Polish children at the prepubertal age. Anthropol. Anz., Aug 17. doi: 10.1127/anthranz/2016/0632.

Sitek, A., Żądzińska, E., Rosset, I., 2012. Effects of psychological stress on skin and hair pigmentation in Polish adolescents. Anthropological Review 75, 1–17.

Sitek, A., Żądzińska, E., Rosset, I., Antoszewski, B., 2013. Is increased constitutive skin and hair pigmentation an early sign of puberty? Homo 64, 205–214.

Slominski, A., Tobin, D.J., Shibahara, S., Wortsman, J., 2004. Melanin pigmentation in mammalian skin and its hormonal regulation. Physiol Rev 84, 1155–1228.

Strapagiel, D., Sobalska-Kwapis, M., Słomka, M., Marciniak, B., 2016. Biobank Lodz - DNA based biobank at the University of Lodz, Poland, Open Journal of Bioresources, in press.

Sturm, R.A., Duffy, D.L., Box, N.F., Newton, R.A., Shepherd, A.G., Chen, W., Marks, L.H., Leonard, J.H., Martin, N.G., 2003. Genetic association and cellular function of MC1R variant alleles in human pigmentation. Ann N Y Acad Sci 994, 348–358.

Sulem, P., Gudbjartsson, D.F., Stacey, S.N., Helgason, A., Rafnar, T., Jakobsdottir, M., Steinberg, S., Gudjonsson, S.A., Palsson, A., Thorleifsson, G., Pálsson, S., Sigurgeirsson, B., Thorisdottir, K., Ragnarsson, R., Benediktsdottir, K.R., Aben, K.K., Vermeulen, S.H., Goldstein, A.M., Tucker, M.A., Kiemeney, L.A., Olafsson, J.H., Gulcher, J., Kong, A., Thorsteinsdottir, U., Stefansson, K., 2008. Two newly identified genetic determinants of pigmentation in Europeans. Nat Genet 40, 835–837.

Sulem, P., Gudbjartsson, D.F., Stacey, S.N., Helgason, A., Rafnar, T., Magnusson, K.P., Manolescu, A., Karason, A., Palsson, A., Thorleifsson, G., Jakobsdottir, M., Steinberg, S., Pálsson, S., Jonasson, F., Sigurgeirsson, B., Thorisdottir, K., Ragnarsson, R., Benediktsdottir, K.R., Aben, K.K., Kiemeney, L.A., Olafsson, J.H., Gulcher, J., Kong, A., Thorsteinsdottir, U., Stefansson, K., 2007. Genetic determinants of hair, eye and skin pigmentation in Europeans. Nat Genet 39, 1443–1452.

Tobin, D.J., 2008. Human hair pigmentation – biological aspects. Int J Cosmet Sci 30, 233–257.

Untergasser, A., Cutcutache, I., Koressaar, T., Ye, J., Faircloth, B.C., Remm, M., Rozen, S.G., 2012. Primer3 – new capabilities and interfaces. Nucleic Acids Res 40, e115.

Valenzuela, R.K., Henderson, M.S., Walsh, M.H., Garrison, N.A., Kelch, J.T., Cohen-Barak, O., Erickson, D.T., John, Meaney F., Bruce, Walsh J., Cheng, K.C., Ito, S., Wakamatsu, K., Frudakis, T., Thomas, M., Brilliant, M.H., 2010. Predicting phenotype from genotype: normal pigmentation. J Forensic Sci 55, 315–322.

Valverde, P., Healy, E., Jackson, I., Rees, J.L., Thody, A.J., 1995. Variants of the melanocyte-stimulating hormone receptor gene are associated with red hair and fair skin in humans. Nat Genet 11, 328–330.

van Beek, N., Bodo, E., Kromminga, A., Gaspar, E., Meyer, K., Zmijewski, M.A., Slominski, A., Wenzel, B.E., Paus, R., 2008. Thyroid hormones directly alter human hair follicle functions: anagen prolongation and stimulation of both hair matrix keratinocyte proliferation and hair pigmentation. J Clin Endocrinol Metab 93, 4381–4388.

Walsh, S., Liu, F., Wollstein, A., Kovatsi, L., Ralf, A., Kosiniak-Kamysz, A., Branicki, W., Kayser, M., 2013. The HIrisPlex system for simultaneous prediction of hair and eye colour from DNA. Forensic Sci Int Genet 7, 98–115.

